# Astrocytes in the retinorecipient superior colliculus display unique cellular and structural properties

**DOI:** 10.1101/2022.08.11.503419

**Authors:** Josien Visser, Jérôme Ribot, Alberto Pauletti, David Mazaud, Christian Henneberger, Nathalie Rouach

**Affiliations:** Neuroglial Interactions in Cerebral Physiology and Pathologies, Center for Interdisciplinary Research in Biology, Collège de France CNRS, INSERM, Labex Memolife, Université PSL, 75005 Paris, France; Doctoral School N°158, Pierre and Marie Curie University, Paris, France; Institute of Cellular Neurosciences, Medical Faculty, University of Bonn, Bonn, Germany; German Center for Neurodegenerative Diseases (DZNE), Bonn, Germany

**Keywords:** superior colliculus, astrocyte, neuroglial interactions

## Abstract

Astrocytes have long been considered to be a largely homogeneous cell population. Recent studies however suggest that astrocytes are highly adapted to the local neuronal circuitry. Glucose utilization in the retinorecipient superior colliculus (SC) is one of the highest in the brain. Since metabolic support to neurons is a major function of astrocytes, they could be of particular relevance in this region and display specific features. However, little is known about astrocytes and their interactions with neurons in this multisensory brain area. We thus here investigated region-specific cellular and structural properties of astrocytes in the visual layer of the SC. Using morphological reconstructions, fluorescent recovery after photobleaching (FRAP) and superresolution imaging, we found that astrocytes from the visual layers of the SC are highly distinct with a higher cellular density, a more complex morphology and a stronger proximity to synapses compared to astrocytes from the primary visual cortex and the hippocampus. These data point to astroglial diversity and specialization within neural circuits integrating sensory information in the adult brain.

## Introduction

Astrocytes can contact thousands of synapses, which they regulate functionally. Hence, the emerging view of the brain is that its functioning arises from orchestrated activity of networks composed of both neurons and astrocytes^1,2^. While neuronal heterogeneity between brain areas has long been appreciated, astrocytes have traditionally been considered a homogeneous cell population across the whole brain. However, recent studies on glial cells revealed that astrocytes can be highly diverse^3–5^. This heterogeneity suggests that astrocytes can have specializations, just like there are different classes of neurons, which significantly changes the way neuron-glia interactions are considered.

Astrocyte heterogeneity is reflected in the highly diverse transcriptomes between brain regions^5,6^, leading to region-specific morphological and functional characteristics, such as calcium signalling^5^, electrophysiological properties and structural features and synapse coverage^7,8^. Another typical feature of astrocytes is their extensive network organization via gap-junction channels. Interestingly, the extent of intercellular coupling and its orientation is region-dependent. In the hippocampus, astrocytic networks from the pyramidal layer are shaped in a columnar manner^9,10^. Astroglial networks are also confined within a barrel field^9,10^. This suggests that these networks are superimposed onto regional neuronal circuits, underlining the adaptive nature of astrocytes to neuronal circuits.

Astrocytic networks play a key role in brain homeostasis and metabolism, for instance by trafficking glucose intercellularly in a neuronal activity-dependent manner^11^. This indicates that astroglial networks functionally adapt to the needs of local neuronal circuits. One of the highest metabolic structures in the brain is the most superficial part of the superior colliculus (SC), where the layers involved in visual processing are located^12,13^. In addition, astroglial glutamate transport mediated by GLT1 contributes to metabolic responses evoked by visual stimulation in the SC^14^, suggesting that astrocytes could be particularly important in this region. The SC is a region in the mammalian midbrain that transforms multisensory information into orienting and approach behaviours. Interestingly, the SC is evolutionary highly conserved, which is reflected in the layered pattern that is present across species. Even neuronal architecture and responses in the SC visual layers are conserved along evolution^15^. However, little is known about astrocytes in the retinorecipient SC.

In this study, we examined region-specific cellular and structural properties of astrocytes in the superficial SC by comparing them to astroglial characteristics from the primary visual cortex (V1) and hippocampus, which are the most extensively studied areas in visual and glia research, respectively.

## Results

### High astrocyte density in the visual layer of the superior colliculus

Neuron and astrocyte densities vary across the brain^16^. We thus compared cellular densities of neurons and astrocytes in the superficial SC, primary visual cortex (V1) and the CA1 region of the hippocampus, using immunostainings for NeuN and Sox9 to label neurons and astrocytes, respectively. In addition, we used a myelin marker, myelin basic protein (MBP), to define the structure subregions (Figure 1a). We found no difference in neuronal densities between the visual layer of the SC and V1 (Figure 1b, SC: n = 12, V1: n = 9, *p* = 0.1226, Tukey’s post hoc test after one-way ANOVA). However, neurons in the hippocampal CA1 region were half as densely populated compared to SC and V1 (CA1: n = 10, SC: n = 12, V1: n = 9, *p* < 0.0001, between SC and CA1 and between V1 and CA1, Tukey’s post hoc test after one-way ANOVA). Furthermore, the astrocytic density was twice larger in the visual layer of the SC compared to the other two brain regions (Figure 1c, CA1: n = 10, SC: n = 12, V1: n = 9, *p* < 0.0001 between SC and V1 and between SC and CA1, Tukey’s post hoc test after one-way ANOVA). As a result, the astrocyte/neuron density ratio was significantly higher in the superficial SC than in V1 (Figure 1d, SC: n = 12, V1: n = 9, *p* < 0.0001, Tukey’s post hoc test after one-way ANOVA). Due to the low neuronal density, this ratio was higher in CA1 compared to SC and V1 (Figure 1d, CA1: n = 10, SC: n = 12, V1: n = 9, *p* < 0.0001 between CA1 and V1 and between CA1 and SC, Tukey’s post hoc test after one-way ANOVA). These results indicate that astrocyte density is particularly high in the visual layer of the SC.

**Figure 1.**
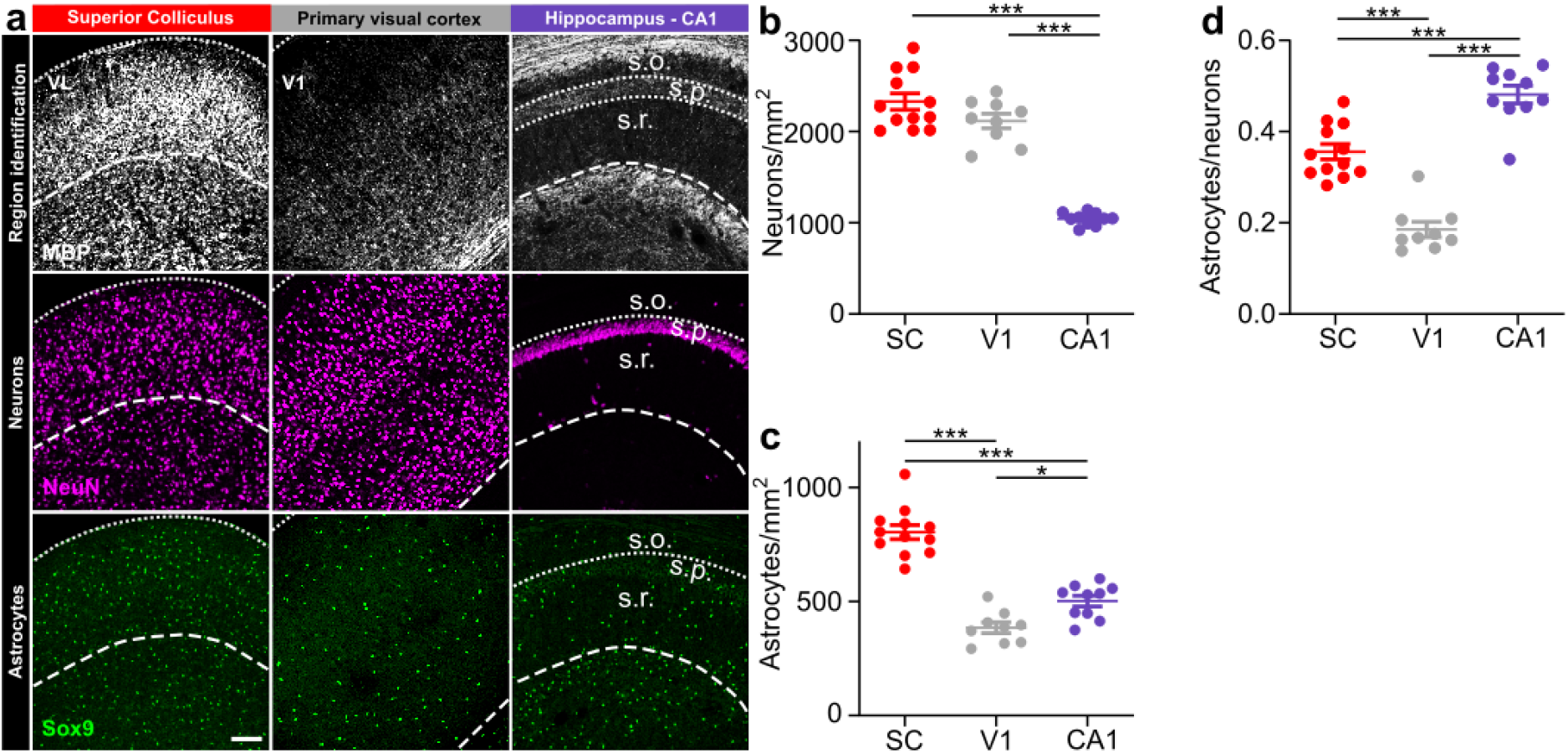
High astrocyte density in the visual layer of the superior colliculus. **(a)** Coronal sections of the visual layer (VL) of the superior colliculus (SC, left panel), primary visual cortex (V1, middle panel) and hippocampal CA1 (CA1, right panel). Regions were identified with a myelin basic protein staining (MBP, top panels). For the hippocampus, the stratum oriens (s.o.), stratum pyrimidale (s.p.) and stratum radiatum (s.r.) are indicated. Representative images for neuron (middle panels) and astrocyte densities (bottom panels). **(b, c)** Quantification of neuron (b) and astrocyte (c) density in the SC (n=12), V1 (n=9) and CA1 (n=10). One-way ANOVA (neuron: F(2,28) = 87.75), *p* < 0.0001, with Tukey’s multiple comparisons \*\*\**p <* 0.0001); astrocyte: F(2,28) = 67.74, p<0.0001, with Tukey’s multiple comparisons \**p* < 0.0001, \*\*\**p* < 0.0001). **(d)** Quantification of astrocyte to neuron density ratio. One-way ANOVA (F(2,28) = 64.15, p < 0.0001, with Tukey’s multiple comparisons \*\*\**p* < 0.0001). n represents the number of brain coronal slices that were analyzed from 3 mice. Scale bar, 25µm. n=3 mice; data are shown as mean ± SEM.

We then investigated whether the density of neurons and astrocytes was heterogenous within the different layers of the SC (Supplementary Figure S1a, b). Densities of both neurons and astrocytes were higher in the visual layer compared to the intermediate and lower layers of the SC (Figure S1b, neurons: n = 12, *p* = 0.0002 between visual and intermediate layers, *p* = 0.0040 between visual and lower layers, Tukey’s post hoc test after repeated measures one-way ANOVA; Figure S1c, astrocytes: n = 12, *p* = 0.0064 between visual and intermediate layers, *p* = 0.0042 between visual and lower layers (n = 12), Tukey’s post hoc test after repeated measures one-way ANOVA).

### Astrocytes in the superior colliculus have a large territory and complex morphology

To assess region-specific morphological characteristics of astrocytes, we analysed the domain area and complexity of GFP-positive astrocytes from superficial SC (n = 8), V1 (n = 11) and hippocampal CA1 (n = 11) of GFAP-eGFP mice. Reconstruction of single astrocytes revealed morphological differences between brain regions (Figure 2a, b). We found no significant differences in soma size between the three brain structures (Figure 2c, *p* = 0.5940, one-way ANOVA). However, astrocytes in the SC cover a larger territory compared to V1 (Figure 2d, *p* = 0.0227 between SC (n = 9) and V1 (n = 11), Tukey’s post hoc test after one-way ANOVA). These data suggest that astroglial processes in the SC may be longer or more numerous to cover a larger area. Therefore, we analysed the arrangement of astrocytic processes (Figure 2e-g), which firstly revealed that the number of processes was higher in the SC compared to V1 (Figure 2e, SC: n *=* 9, V1: n = 11, *p* = 0.0140, Tukey’s post hoc test after one-way ANOVA). This increase was caused by more short processes with length between 5 and 15 µm (Figure 2g, SC: n *=* 9, V1: n = 11, CA1: n = 11, 10-15 µm: *p* < 0.0001 between SC and V1 and, *p* = 0.0311 between SC and CA1; 10-15µm: *p* = 0.0034 between SC and V1, *p* = 0.0056 between SC and CA1, Tukey’s post hoc test after two-way ANOVA). Altogether, this indicates that astrocyte territory is increased in the SC compared to V1 through a larger number and an increased length of short processes. To test this hypothesis, we performed Sholl analysis, which revealed that collicular astrocytes display a more complex organization close to the soma compared to both V1 and CA1 (Figure 2j, 5-10 µm, *p* = 0.0012 between SC (n = 9) and V1 (n = 11), *p* = 0.0055 between SC and CA1 (n = 11); 10-15 µm: *p* < 0.0001 between SC and V1 and between SC and CA1; 15-20 µm: *p* = 0.0002 between SC and V1, *p* = 0.0066 between SC and CA1, Tukey’s post hoc test after two-way ANOVA). This data thus shows that astrocytes in the visual layer of the SC have more proximal complexity compared to V1 and CA1.

**Figure 2.**
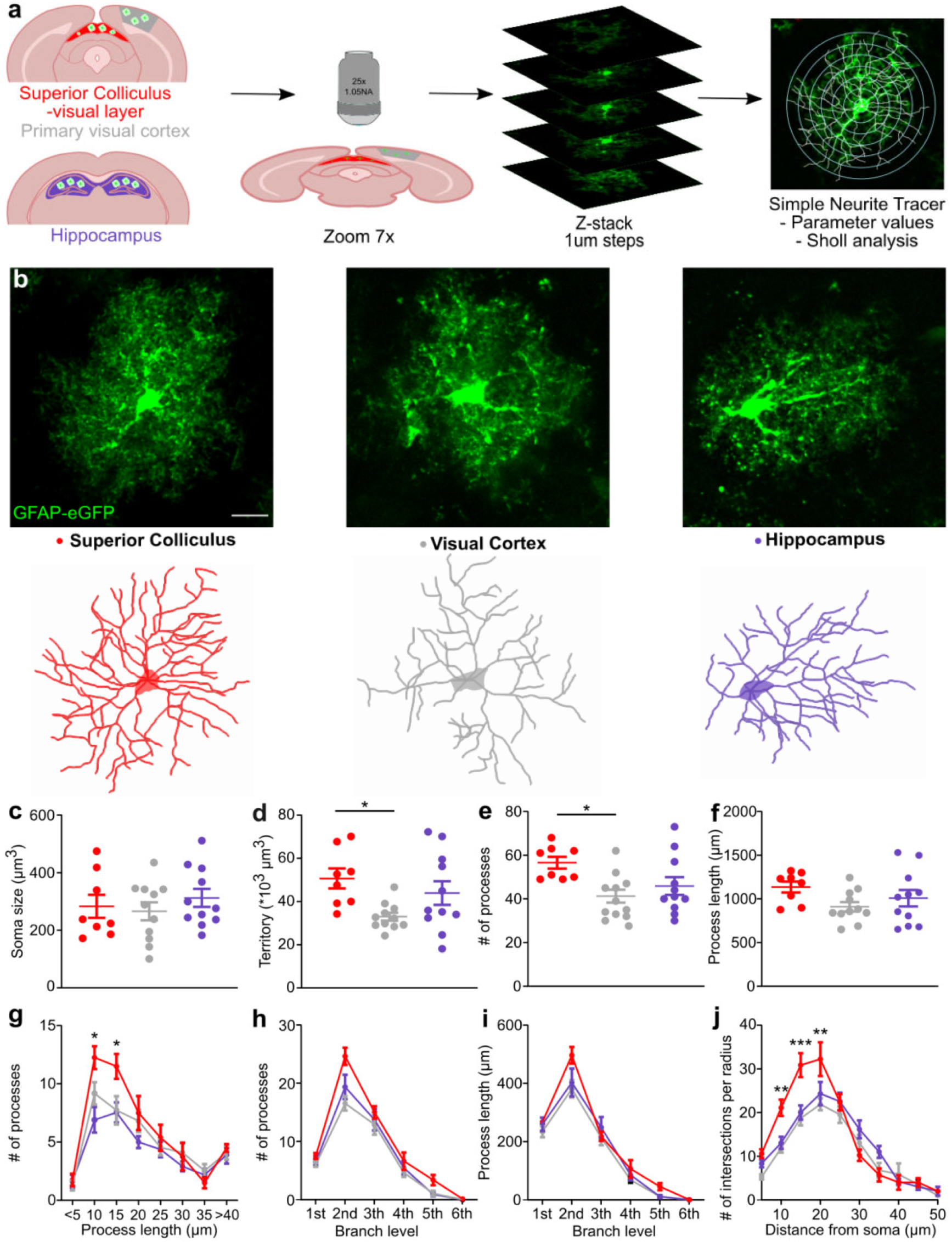
Astrocytes in the visual layer of the superior colliculus are large and complex. **(a)** Astrocyte morphology in the visual layer of the superior colliculus (SC, red), primary visual cortex (V1, grey) and hippocampal CA1 region (CA1, purple) in acute brain slices from GFAP-eGFP mice. **(b)** Representative images of astrocytes in the areas of interest (top panels) and their corresponding reconstructions (bottom panels). **(c)** Quantification of the soma size in the three regions (One-way ANOVA, F(2,27) = 0.5310, *p* = 0.5940, with Tukey’s multiple comparisons). **(d)** Quantification of astrocyte territories (One-way ANOVA, F(2,27) = 4.252, *p* = 0.0248, with Tukey’s multiple comparisons \**p* < 0.05). **(e-f)** Quantification of number of processes (e) and process length (f) in SC, V1 and CA1 (One-way ANOVA, number of processes: F(2,27) = 4.711, *p* = 0.0176 with Tukey’s multiple comparisons \**p* < 0.05); process length: F(2,27) = 2.121, *p* = 0.1395) **(g)** Quantification of processes per process length (Two-way ANOVA, F(14, 216) = 1.716, *p* = 0.0460 with Tukey’s multiple comparisons \**p* < 0.01), **(h)** number of processes per branch level (Two-way ANOVA, F(10, 162) = 1.048, *p* = 0.4058 and **(i)** total process length per branch level (Two-way ANOVA, F(10, 162) = 1.468, *p* = 0.1557. **(j)** Quantification of astrocyte complexity by Sholl analysis (Two-way ANOVA, F(18, 217) = 2.883, *p* = 0.0001 with Tukey’s multiple comparisons \*\**p* < 0.001, \*\*\**p* < 0.0001). Scale bar, 10 µm. SC: n = 8 astrocytes, V1: n = 11 astrocytes, CA1: n = 11 astrocytes. n represents the number of cells analysed from 3 mice. Data are shown as mean ± SEM.

### Low intracellular diffusion in astrocytes from the superficial superior colliculus

We then determined whether the high morphological complexity of astrocytes in the visual layer of the SC translated functionally into an alteration in the intracellular diffusion properties of individual astrocytes. To do so, we used the FRAP technique in acute brain slices from GFAP-eGFP mice, where the gap-junction impermeable cytosolic eGFP expressed by astrocytes was bleached by performing line scanning experiments at an increased laser intensity (Figure 3a). Closure of the laser shutter led to fluorescence recovery, because it allows intact eGFP to diffuse into the bleached volume. FRAP was analysed as illustrated (Figure 3a) to measure cytosolic diffusivity. We found that FRAP was about half in whole astrocytes from the SC compared to astrocytes from V1 and CA1 (Figure 3b, *p* = 0.0170 between SC (n = 8) and V1 (n = 10), *p* = 0.0004 between SC and CA1, one-way ANOVA followed by Tukey’s post hoc test). These region-specific differences did not involve changes in somatic diffusivity (Figure 3c, SC: n = 7, one value was excluded as it was negative, V1: n = 10, CA1: n = 10, *p* = 0.4528, one-way ANOVA), but originated from reduced intracellular diffusion in astroglial processes (Figure 3d, *p* = 0.0167 between SC (n = 8) and V1 (n = 10), *p* = 0.0004 between SC and CA1 (n = 10), Tukey’s post hoc test after one-way ANOVA). These data indicate that the high morphological complexity of astrocytes from the visual layer of the SC is accompanied by a lower intracellular diffusivity specifically in the processes.

**Figure 3.**
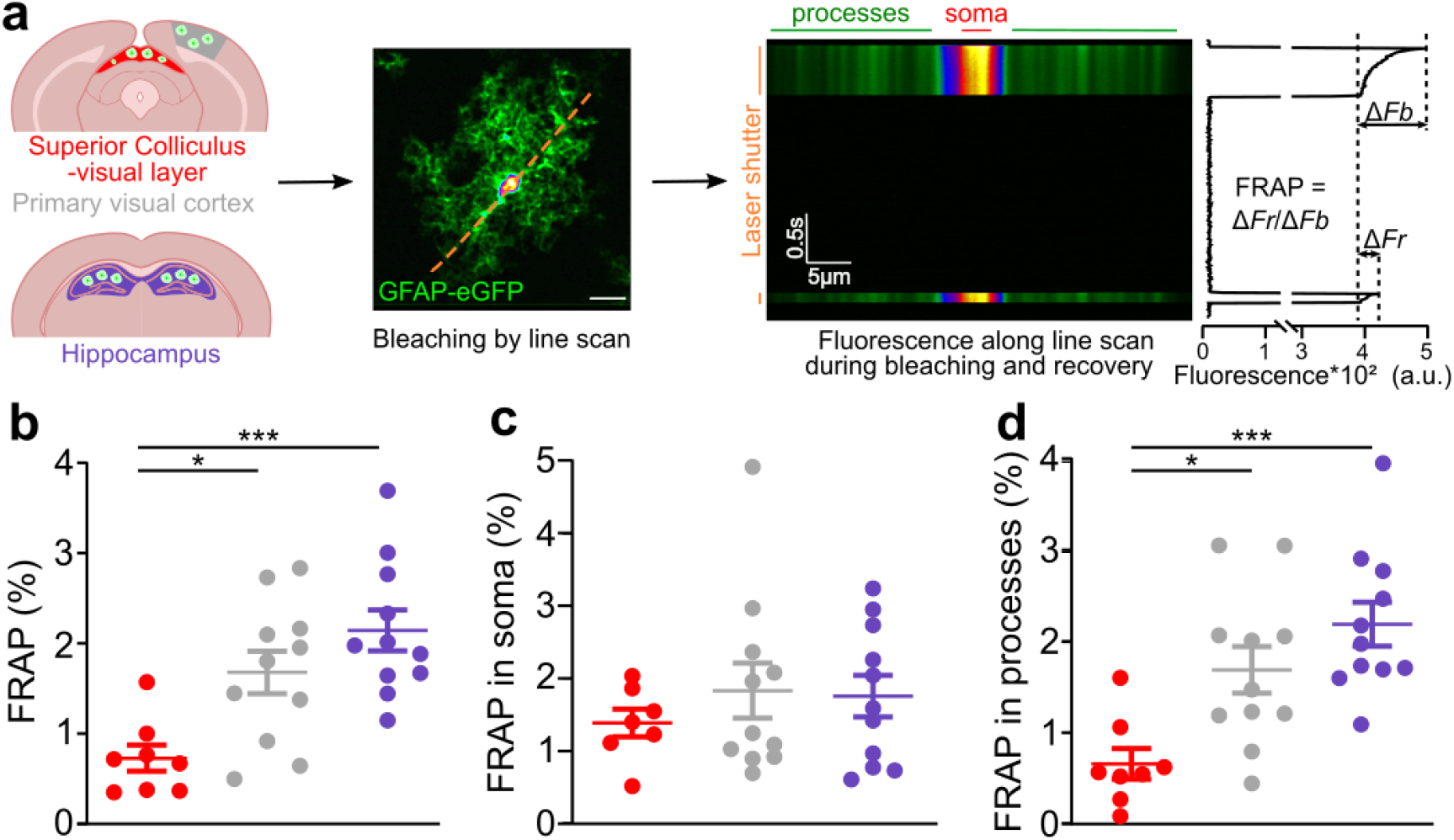
Low intracellular diffusion in astrocytes from the visual layer of the superior colliculus. **(a)** Schematic overview showing the approach used to quantify intracellular diffusion in astrocytes from different brain areas, the visual layer of the superior colliculus (SC), the primary visual cortex (V1) and the CA1 region of the hippocampus (CA1). Cytoplasmic eGFP expressed by astrocytes was bleached by high laser power line scanning. Recovery occurred when unbleached GFP entered the scanned area. Fluorescence Recovery After Photobleaching (FRAP) was calculated by dividing the bleaching factor (Δ*Fb*) by the recovery fraction (Δ*Fr*). **(b)** FRAP was significantly reduced in the SC (n =8) compared to V1 (n = 11) and CA1 (n = 11) (One-way ANOVA, F(2,27) = 9.806, *p* = 0.0006, with Tukey’s multiple comparisons \**p* < 0.05, \*\*\**p* < 0.001). (**c**) Diffusion speed in the soma was similar (One-way ANOVA, F(2,26) = 0.4528, *p* = 0.6407). (**d**) The difference in intracellular diffusion speed arises from the processes, as FRAP is lower in the SC (One-way ANOVA, F(2,27) = 9.840, *p* = 0.0006, with Tukey’s multiple comparisons \**p* < 0.05, \*\*\**p <* 0.0001). Scale bar, 10 µm. n represents the number of cells analysed from 3 mice. Data are shown as mean ± SEM.

### Stronger coverage of synapses in astrocytes from the visual layer of the superior colliculus

Astrocytes are integrated components of the tripartite synapse, which is composed of a presynaptic and postsynaptic element together with an astroglial process. The tight structural interactions between these elements allow astrocytes to actively regulate synapse function (Figure 4a, upper left panel)^17^. Since astroglial processes from the visual layer of the SC display a high complexity, we investigated whether this resulted in tighter fine structural interactions between astroglial processes and synapses. To visualize and quantify structural properties of tripartite synapses in the visual layer of the SC, we used super-resolution STED microscopy (Figure 4a, lower left panel)^18^ and measured the closest distance between synapses and astrocytic processes from GFAP-eGFP mice (Figure 4a, right panel). We found that astrocytic processes from the superficial SC were closer to synapses compared to the ones from the CA1 or V1 region (Figure 4b-c, *p* < 0.0001 between SC (n = 6710 synapses) and CA1 (n = 9837 synapses), *p =*0.001 between SC and V1 (n = 8702 synapses), Kolmogorov-Smirnov test). Further analysis of the distance distributions showed that the visual layer of the SC displayed a higher proportion of synapses in close proximity (< 210nm) to astrocytic processes compared to V1 and hippocampal CA1 (Figure 4d and 4e, *p* < 0.0001 between SC (n = 16 astrocytes) and V1 (n = 17 astrocytes) and between SC (n = 16 astrocytes) and CA1 (n = 13 astrocytes), Tukey’s post hoc test after one-way ANOVA).

**Figure 4.**
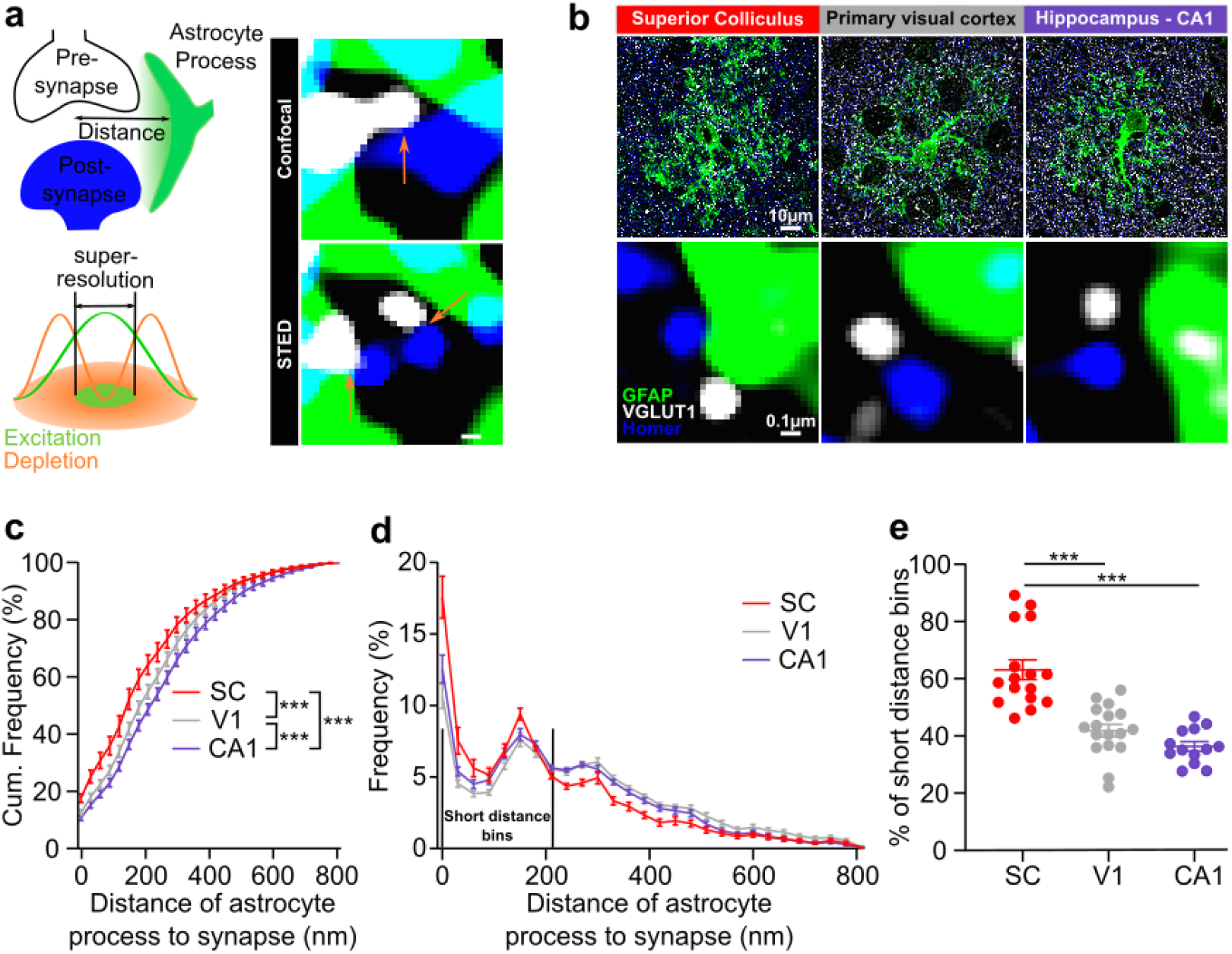
Tight astroglial coverage of synapses in the visual layer of the superior colliculus. **(a)** Schematic illustrating the method used to measure the distances between astrocytic processes and synapses (left, top panel) with super-resolution STED microscopy combining an excitation and a depletion laser (left, bottom panel). Using this method, synaptic boutons can be accurately distinguished (right, lower panel), as shown in the representative images compared to confocal microscopy (right, top panel). Scale bar, 0.1µm. **(b)** Top: representative confocal images of astrocytes labelled with eGFP in the visual layer of the superior colliculus (SC), primary visual cortex (V1) and hippocampal CA1 (CA1), together with VGLUT1 and Homer to visualize pre- and postsynaptic elements, respectively. Scale bar, 10µm. Bottom: representative STED images of an astrocytic process with a synapse in each brain structure. Scale bar, 0.1 µm. **(c)** Cumulative frequency of the distances between astrocyte process and synapse. Kolmogorov-Smirnov (SC (n = 6710 synapses) vs V1 (n = 8702 synapses): \*\*\**p* < 0.0001; SC vs CA1 (n = 9837 synapses): \*\*\**p* < 0.0001; V1 vs CA1: \*\*\**p* < 0.0001). **(d)** Frequency distributions of distance between processes and synapses. Short distances were defined as distances up to 210 nm. **(e)** Quantification of the number of short distance processes (SC: n = 16, V1: n = 17, CA1: n = 13, One-way ANOVA, F(2,43) = 28.45, *p* < 0.0001, with Tukey’s multiple comparisons \*\*\**p <* 0.0001). n is represented as number of astrocytes, which are acquired from 4 mice, unless differently indicated, data are shown as mean ± SEM.

We then investigated whether the increased astroglial coverage of synapses in the superficial SC could result from an increased synapse density along astroglial processes (Figure S2a). Using STED microscopy, we however found that the synapse density per volume of astrocytic process was only particularly high in the hippocampal CA1 region compared to the visual SC and V1 (Figure S2b, *p* < 0.0001 between CA1 (n = 13) and SC (n = 16 astrocytes) and between CA1 and V1 (n = 17 astrocytes), Tukey’s post hoc test after one-way ANOVA). In all, this indicates that the spatial arrangement of the tripartite synapses is region-specific and that the astroglial processes-synapses structural interactions are tighter in the visual layer of the SC.

## Discussion

While neuronal heterogeneity has long been established, until recently astrocytes have been considered a homogeneous ‘glue’ filling the space between neurons. However, recent studies point to astrocytes as highly diverse^19–21^ cells and specialized to neuronal circuits^7,22^. Moreover, astrocytes play a vital role in brain metabolism and are directly metabolically coupled to neurons^23^. Here, we studied region-specific properties of astrocytes in the SC, one of the highest metabolic structures in the brain^12^ involved in multisensory processing. We reveal that astrocytes in the SC indeed differ from astrocytes in V1 and hippocampus at the structural and population levels, indicating that astrocyte heterogeneity can be observed at different scales.

At the population level, astrocytic cellular densities are higher in the visual layer of the SC compared to hippocampal CA1 and V1. Since astrocytes support the energetic need of neurons within their territory^23^, it could be hypothesized that in the visual SC, more astrocytes are necessary to account for the high metabolic cost within this area^13,24^.

At the single cell level, we observed that astrocytes in the visual layer of the SC occupy a larger territory compared to the ones in another visual structure (V1). Astrocytes are known for their highly complex morphologies, which are in contact with synapses^17^, blood vessels^25^ and other glial cells^11^. Hence, we compared astroglia morphology between our three areas of interest. We observed that astrocytes within the visual SC are more complex close to the soma and have more shorter processes. Using 2-photon imaging of ex-vivo brain tissues, we revealed the presence of more complex astroglial processes primarily at the proximal level. These morphological differences translate functionally into a slow intracellular diffusion within astroglial processes.

Astrocytes are known to change their morphology and coverage of synapses in response to neuronal activity^26^ and can thereby actively regulate synaptic transmission^27–29^ and plasticity^30^. Hence, after determining single astrocyte morphology, we investigated potential differences at the tripartite synapse level by determining the astroglial coverage of synapses. In the visual SC, the astrocytic processes are closer to excitatory synapses compared to the hippocampus and visual cortex. One of the functions of astroglial perisynaptic processes is to provide neurons with energy substrates, for example through the astrocyte-neuron lactate shuttle^31^. Hence, these closer astroglial processes to synapses in the visual SC could quickly provide neurons with lactate upon neuronal activation. Also, glutamate uptake could be more efficient in the SC, and extracellular spread of synaptically-released glutamate more limited, because synaptic astrocyte coverage correlates with the efficacy of glutamate uptake^29,32^ and acute withdrawal of perisynaptic astrocytic processes increases glutamatergic synaptic crosstalk as shown for the hippocampus^28^.

In summary, in this study, using a variety of approaches in the adult mouse, we investigated whether astrocytes in the visual SC have region-specific features. We revealed that astrocytes in the visual SC are highly different at the population, whole-cell, intracellular and even at the nanoscale tripartite synapse levels. Future studies could elucidate whether these region-specific characteristics of astrocytes in the visual SC have implications for the functioning of synapses, neurons and neuronal circuits as well as for the high metabolic demand of this brain structure.

## Acknowledgements

The authors thank Philippe Mailly from the imaging platform for the analysis of STED images.

## Methods

### Animals

All experiments were performed in accordance with the European Communities Council Directives of 01/01/2013 (2010/63/EU) for animal care and experimentation and of the French ethic committee (ethics approval #201902121059308 delivered by the French ministry of higher education, research and innovation). Experiments were carried out using mice of wildtype C57BL/6j background (Janvier labs, France), and mice expressing the enhanced green fluorescent protein under the astroglial promoters glial fibrillary acidic protein (GFAP-eGFP)^33^, which were obtained from F. Kirchhoff (University of Saarland, Germany). Adult mice of both genders were used at postnatal days 50 to 100. All mice were housed under standard conditions (12-hour light/ 12-hour dark cycle, light on at 7 am, 22±1°C ambient temperature, 60% relative humidity), with ad libitum access to food and water. All efforts were made to minimize the number of animals used and their suffering.

### Astrocyte morphology

After rapid extraction of GFAP-eGFP mouse brains, coronal slices with 350 µm thickness containing the SC, V1 or hippocampus were cut using a vibratome (Leica VT1200S) in ice-cold aCSF composed of (in mM): 119 NaCl, 2.5 KCl, 2.5 CaCl_2_, 1.3 MgSO_4_, 1 NaH_2_PO_4_, 26.2 NaHCO_3_ and 11 glucose. Slices were allowed to recover for a minimum of 30 min in a chamber containing aCSF at room temperature. In all experiments, aCSF was continuously bubbled with 95% O_2_/5% CO_2_. Slices were then placed into a submersion-type recording chamber superfused with oxygenated aCSF at 34°C on a 2-photon excitation fluorescence microscope (FV10MP, Olympus) linked to a femtosecond Ti:sapphire 80 MHz pulse laser (Vision S, Coherent). Z-stacks of whole astrocytes in the superficial SC, V1 and hippocampal CA1 were acquired with a 20x objective with 7x zoom at 1µm intervals using a 800 nm laser.

Images were analysed using Simple Neurite Tracer (SNT), an open-source plugin developed for ImageJ^35^, that enables semi-automated morphological reconstruction and Sholl analysis of astrocytes^36^. After performing tracing of astrocytic processes through the whole cells, Sholl analysis was performed, counting the number of radial crossings with concentric circles at 5 µm intervals. Additionally, SNT enables the quantification of the number of processes and process length (Figure 2a). Investigators were blinded to the morphological analysis.

To define astrocytic territory and soma size, background subtraction and a median filter in ImageJ were applied followed by Otsu-thresholding. Close- and filling holes functions were performed on the image. Afterwards, a macro was used to measure the volume of the thresholded area for each image in the stack for the soma as well as the whole astrocytic territory.

### Fluorescence Recovery After Photobleaching (FRAP)

After acquiring images for morphological analysis, the FRAP technique was applied to study intracellular diffusivity in single astrocytes^37^. Bleaching of the eGFP-signal was induced by increasing the laser power to 15-30mW for 500 ms and performing a line scan, which comprised both the soma and astrocytic processes. Afterwards, eGFP fluorescence recovery was allowed by closing the laser shutter for 1 s, before line scanning was resumed. The bleached fluorescence (Δ*Fb*) and the recovered fluorescence (Δ*Fr*) were determined and used to calculate FRAP (FRAP = Δ*Fr/* Δ*Fb*) of the whole astrocyte and also separately for astrocytic processes and soma (Figure 3a).

### Slice preparation and Reagents for immunohistochemistry

C57Bl6/6j mice were anesthetized with a lethal dose of Euthasol (150 mg/kg, Intraperitoneal injection), and transcardially perfused with phosphate buffered saline (PBS) followed by 2% paraformaldehyde (PFA) in PBS. After perfusion, brains were carefully dissected and postfixed for 24hours in 2% PFA followed by 24 hours in 30% sucrose solution for cryoprotection. Brain coronal sections (40 µm) of slices containing the hippocampus or both the SC and V1 were cut with a Leica microtome and stored in PBS. Free floating slices of the hippocampus, SC and V1 were incubated 2 hours with PBS-1% gelatine with 0.25% Triton-X100 (PGT) to block unspecific binding sites.

For confocal imaging, slices from C57Bl/6j mice were prepared as follows: brain sections were incubated at 4°C with the following primary antibodies: MBP (rabbit, 1:300, Novus Biologicals), Sox9 (goat, 1:500, AF3075, R&D systems) and NeuN (mouse, 1:500, Sigma). 24 hours later the slices were washed three times in PGT following by an incubation of 2 hours on room-temperature with the following secondary antibodies: anti-rabbit 488 (donkey, 1:1000, Life Technologies), anti-goat 568 (donkey, 1:1000, ThermoFisher) and anti-mouse 647 (donkey, 1:1000, Life Technologies). Finally, the slices were washed three times in PBS and mounted with Fluoromount-G (Southern Biotechnology).

For STED imaging, slices from GFAP-eGFP mice were incubated at 4°C for 72 hours with anti-GFP (chicken, 1:500, Aves 1020), anti-VGLuT1 (mouse, 1:200, Synaptic Systems, #135511) and anti-Homer-1 (rabbit, 1:200, Synaptic Systems, #160003) as primary antibodies. After three washes with PGT, the following secondary anitbodies were applied for 72 hours: anti-Chicken Alexa (goat, 1:300, Invitrogen, A11039), anti-Rabbit Alexa594 (goat, 1:200, Jackson ImmunoResearch, # 111-586-045) and anti-Mouse Alexa647 (goat, 1/200, Life Technologies, #A21235). Lastly, slices were washed three times in PBS and mounted with solid antifade mounting medium (Abberior).

### Confocal and STED Imaging and analysis for immunohistochemistry

Confocal imaging was performed using an inverted confocal laser-scanning microscope (Confocal Leica SP5 inverted). Images were taken with a 20x objective and acquired sequentially with 543 nm and 647 nm lasers. For each animal, three images were recorded per brain region (hippocampal CA1, V1 and superficial SC). The images were then analysed using ImageJ. First, areas were determined using MBP (Figure 1a). For the Sox9 and NeuN signals, background subtraction was applied as well as a gaussian blur. To determine neuronal and astrocytic densities, the local maxima over the images were detected (Astrocytes: prominence > 150, Neurons: prominence > 80).

STED images were taken using an upright STED microscope (Abberior Instruments GmbH) previously described^38,39^. Briefly, this STED-microscope has 488nm, 561nm and 640nm confocal excitation lasers (Abberior Instruments, pulsed @40/80 Mhz) together with a STED-laser of 775nm (MPB-C,pulsed @40/80 Mhz) (Figure 4a). For each brain region, one or two astrocytes per slice were imaged. For each astrocyte, 3 to 10 field of views with a z-stack of 0.03µm intervals were acquired. Homer-1 and VGluT1 signals were acquired with STED and confocal resolutions simultaneously, while GFAP signal was only acquired with confocal resolution. These images were then deconvolved using the Huygens software and analysis was performed using the ImageJ software combined with an internally developed plugin to determine the distance from a detected excitatory synapse to the closest astrocytic process^40^. In short, peak intensities of VGlut-1 and Homer-1 were identified in the STED images. To remove false positives, the existence of a signal for these puncta was confirmed in the corresponding deconvolved confocal images. A pair of VGlut-1/Homer-1 punctae was defined as a synapse if these two punctae lay within 300 nm of each other. Finally, the distance between the synapse to the closest astrocyte process was computed with a Li-thresholded^41^ image of the deconvolved astrocyte. The synapse density around astroglial processes was calculated by dividing the number of synapses within the astroglial environment (up to 800 nm) divided by the astroglial volume within that image.

### Statistical analysis

All data are expressed as mean ± standard error of the mean (SEM). Statistical analysis was performed with GraphPad Prism (GraphPad software). Normality of the distributions and the equality of the standard deviations were tested with D’Agostino-Pearson normality and Brown-Forsythe test respectively. Based on these outcomes, appropriate parametric or nonparametric were used.

