## Supplemental Figures for "Astrocytes in the retinorecipient superior colliculus display unique cellular and structural properties"

### Supplementary figures

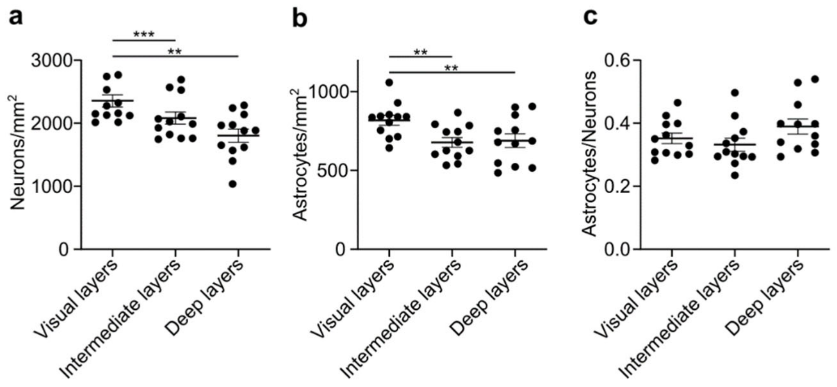


**Figure S1. Higher cellular densities in the visual layers for the superior colliculus (a)** Quantification of astroglial densities in the superficial/ visual layers (n = 12), the intermediate (n = 12) and deep layers (n = 12) of the SC (Repeated measures ANOVA, $F(1.132, 12.45) = 11.26$, $p = 0.0045$, with Tukey's multiple comparisons $^{**}p < 0.001$, $^{***}p < 0.0001$). **(b)** Quantification of neuronal densities in the superficial/ visual layers, the intermediate and deep layers of the SC (Repeated measures ANOVA, $F(1.494, 16.44) = 7.711$, $p = 0.0073$, with Tukey's multiple comparisons $^{**}p < 0.001$). **(c)** Quantification of astrocyte to neuron ratio in the different regions within the SC (Repeated measures ANOVA, $F(1.277, 14.04) = 2.073$, $p = 0.1708$). Data are shown as mean ± SEM. n represents the number of slices

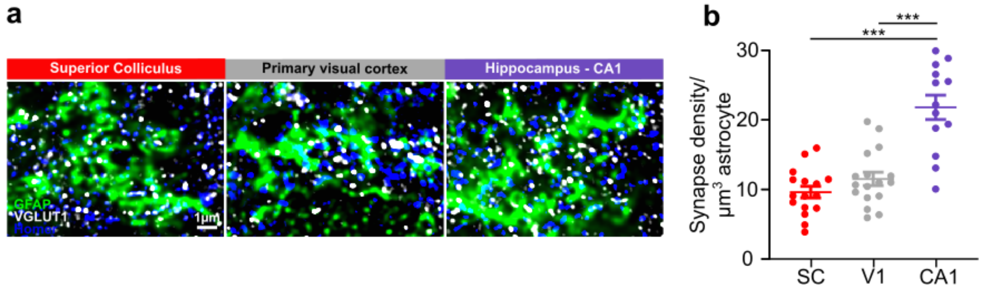


**Figure S2. Synapse density heterogeneity around astroglial processes. (a)** Representative images of an astrocytic process with surrounding synapses in each brain region. **(b)** In hippocampal CA1 there are more synapses per astrocytic volume, indicating that synaptic densities along the astrocytic processes are higher in this area (SC: n = 16, V1: n =17, CA1: n = 13, one-way ANOVA ($F(2,43) = 28.73$, $p < 0.0001$ with Tukey's multiple comparisons $^{***}p < 0.0001$). Data are shown as mean ± SEM. n represents the number of cells analysed from 3 mice.
